# Within-host Evolution of Segments Ratio for the Tripartite Genome of *Alfalfa Mosaic Virus*

**DOI:** 10.1101/066084

**Authors:** Beilei Wu, Mark P. Zwart, Jesús A. Sánchez-Navarro, Santiago F. Elena

**Author notes:** Current address: Institute of Plant Protection, Chinese Academy of Agricultural 9 Sciences, Beijing, China (Beilei Wu). Current address: Institute of Theoretical Physics, University of Cologne, Cologne, 11 Germany (Mark P. Zwart).

## Abstract

One of the most intriguing questions in evolutionary virology is why multipartite viruses exist. Several hypotheses suggest benefits that outweigh the obvious costs associated with encapsidating each genomic segment into a different viral particle: reduced transmission efficiency and segregation of coadapted genes. These putative advantages range from increasing genome size despite high mutation rates (i.e., escaping from Eigen’s paradox), faster replication, more efficient selection resulting from segment reassortment during mixed infections, or enhanced virion stability and cell-to-cell movement. However, empirical support for these hypotheses is scarce. A more recent hypothesis is that segmentation represents a simple and robust mechanism to regulate gene copy number and, thereby, gene expression. According to this hypothesis, the ratio at which different segments exist during infection of individual hosts should represent a stable situation and would respond to the varying necessities of viral components during infection. Here we report the results of experiments designed to test whether an evolutionary stable equilibrium exists for the three RNAs that constitute the genome of *Alfalfa mosaic virus* (AMV). Starting infections with many different combinations of the three segments, we found that, as infection progresses, the abundance of each genome segment always evolves towards a constant ratio. Population genetic analyses show that the segments ratio at this equilibrium is determined by frequency-dependent selection; indeed, it represents an evolutionary stable solution. The replication of RNAs 1 and 2 was coupled and collaborative, whereas the replication of RNA 3 interfered with the replication of the other two. We found that the equilibrium solution is slightly different for the total amounts of RNA produced and encapsidated, suggesting that competition exists between all RNAs during encapsidation. Finally, we found that the observed equilibrium appears to be host-species dependent.

**Author Summary:** This research focuses on the evolution of genome segmentation, the division of an organism’s hereditary material into multiple chromosomes. Why has the genome evolved these partitions? When is it advantageous to divide the genome over multiple segments? In the case of RNA viruses segmentation may provide a robust and yet tunable mechanism to regulate the expression of different genes. To explore this possibility, we used a tri-segmented plant RNA virus and found that, as expected under this hypothesis, during infection the system evolves towards an optimal solution. The solution varies among host plant species, suggesting that genome segmentation allows for the rapid adaptation to different host plant species. Genome partition can therefore be seen as a stable yet readily adaptable manner to regulate expression of virus genes, by means of gene copy-number variation. We proposed a novel, general evolutionary framework to analyze and interpret quantitative data on segments relative abundances.

## Introduction

The highest level of physical organization of the genome is the division of the hereditary material into multiple segments. Genome segmentation is a ubiquitous feature of eukaryotes, with nuclear chromosome numbers covering an enormous range: from 2*n* to 630*n* [1, 2]. In contrast, bacteria and archaea typically have a 64 single chromosome [3]. Although many viruses also have only a single genome segment, in some species the genome has been partitioned in multiple segments [4, 5]. Whereas most viruses package multiple segments into a single virus particle (*e.g*., reovirus, orthomyxovirus), some plant viruses package each segment into a separate virus particle, a property known as multipartition. In the extreme case, nanoviruses have up to eight DNA genome segments plus several satellite-like segments packed up into different viral particles, although not all segments must enter a cell to cause infection [4]. For multipartite RNA viruses, the number of segments is typically lower, ranging from two to five. It is thought that all genome segments must enter the same cell to establish infection [5].

The evolution of segmented genomes revolves around tradeoffs between potential costs and benefits inherent to different genome architectures. An obvious cost of multipartition – the allocation of each genome segment to different virus particles – is the necessity of coinfecting cells with all viral particles to ensure the presence of at least one copy of the less abundant segment, a cost that increases with the number of segments and particles [6]. All else being equal, an equimolecular composition of particles would maximize the probability of initiating the infection of a host cell successfully. Deviations from this situation would increase the cost of multipartition. Another potential cost of genome segmentation would be the breakage of co-adapted groups of genes during coinfection with several strains of the virus [7]. Several advantages have been proposed to compensate for these costs: (*i*) for the high mutation rates of most RNA viruses, smaller segments are more likely to be copied without errors than larger segments [8], (*ii*) smaller genomic segments should be replicated faster [9], (*iii*) segmentation favors genomic reassortment and thus increases genetic variability by rapidly bringing together beneficial mutations that have occurred in different lineages [10, 11], minimizing the effect of clonal interference [12] and speeding up the rate of adaptation, (*iv*) encapsidation of smaller genomes results in enhanced capsid stability [13], (*v*) particularly in the case of plant viruses, smaller capsids would facilitate trafficking throughout the size-limiting plasmodesmata [14], and (*vi*) segmentation represents an efficient yet simple way to control gene expression by regulating gene copy numbers [15].

Starting on the mid-seventies, a number of publications have addressed different aspects of the replication and regulation of gene expression of plant multipartite viruses, specially for members of the *Bromoviridae* family such as *Brome mosaic virus* (BMV) and *Cowpea chlorotic mottle virus* (CCMV). Many interesting conclusions were drawn in these studies, but particularly relevant for the problem of the evolution of multipartite virus are: (*i*) the ratio of RNA segments varies among closely related viral species (BMV and CCMV) [16] and even among different BMV isolates [17].(*ii*) The relative abundances of genomic segments for bromoviruses is host-dependent [18, 19].(*iii*) Mutations in coding and noncoding sequences of different genomic segments have a profound impact on the accumulation of the other segments [20-24].

Sicard et al. [15] monitored the frequency of the eight single-gene-encoding segments, made of circular DNA, that constitute the genome of the nanovirus *Faba bean necrotic stunt virus* (FBNSV) during infection of single host plants. They observed that regardless of the initial ratio of segments in the inocula, the ratio of segments always evolved towards a constant composition that the authors designated as the “setpoint genome formula” (hereafter referred as *SGF*), which did not represent an equimolecular mixture of genomic segments. They also found that the *SGF* corresponds to a state of maximal viral accumulation and of enhanced symptoms, thus suggesting that segmentation has evolved as a mechanism to regulate gene expression. Finally, in agreement with the bromoviruses results, they also found that the exact stoichiometry of the *SGF* depends on the host plant species. FBNSV, although being a well suited model system for addressing questions related to genome segmentation and multipartition, is not very representative of most known multipartite plant viruses: most are RNA viruses and have a lower number of genome segments.

As mentioned above, the molecular biology of multipartite RNA viruses have been extensively studied [16-24], although it is not known for these viruses whether an *SGF* exists and whether it is evolutionarily stable. In other words: we still miss an evolutionary genetic mechanism that explains the evolution of multipartition in RNA virus populations. This omission is surprising, given that the crucial importance of the existence of such genome ratio equilibria. Genome segment ratios will affect gene expression and, together with regulation at the transcriptional level, ultimately determine the development of infection at the cellular and organismal level. If no stable *SGF* exists, genome segment ratios could be in a state of perpetual flux, and some hypotheses for the advantages of segmentation would need to be discarded or revisited. If multiple stable equilibria exist instead of a single *SGF*, a virus could potentially alternate between different gene expression and infection states, without leaving a signature at the nucleotide-sequence level. Similarly, if different *SGF* exist in different host species, this would suggest that segmentation can facilitate adaptation by rapid regulation of gene expression in a mutation-independent manner [15]. The nature of the *SGF* for RNA viruses therefore has broad implications for our understanding of their infection dynamics and evolution.

The aim of our study is threefold. First, we sought to explore whether a *SGF* also exists for a prototypical multipartite RNA virus. Second, we also set out to determine the effect of host species on the stoichiometry of the *SGF*. Third, we also propose a novel analytical and computational framework, of universal applicability, to the evolutionary analysis of the abundance of any number of segments in segmented viral genomes. To tackle these questions, we have chosen *Alfalfa mosaic virus*(AMV; genus *Alfamovirus*, family *Bromoviridae*), whose genome is composed of three positive-sense ssRNA molecules (RNAs 1, 2 and 3). Briefly, RNAs 1 and 2 encode for proteins essential for replication (P1 and P2) while RNA3 encodes for the movement (MP) and coat (CP) proteins, the latter being translated from a subgenomic RNA4 produced by transcription of the negative-sense strand of RNA3 [25]. Our results show that an evolutionary stable but host-species-dependent *SGF* occurs for this prototypical plant RNA virus.

## Results

### Determination of the *SGF* for AMV

To determine the *SGF* for AMV, we performed inoculation experiments with a range of RNAs 1, 2 and 3 ratios.*Nicotiana benthamiana* plants were mechanically inoculated with a constant amount of RNA but varying the proportions of the three segments as detailed in the Methods section. If a *SGF* exists, then we expect that these different combinations will all evolve towards it. As infection progressed, we sampled different tissues at different stages of infection and estimated the abundance of each RNA on the samples by RT-qPCR. Two types of RNA samples were prepared from each tissue: total RNA and encapsidated RNA from purified viral particles. The first sample represents the total amounts of RNAs 1, 2 and 3 synthetized during infection whereas the second sample represents the amount of each RNA that has been encapsidated and thus is expected to be the relevant figure in terms of horizontal virus transmission.Table 1A shows the results of the multivariate analysis of variance (MANOVA) analysis for the RNA frequencies from total RNA extractions fitted to equation (1) in Methods. All factors contribute in a highly significant manner to the observed variability in RNA segments frequency (in 169 all cases *P* < 0.001).

**Table 1.**
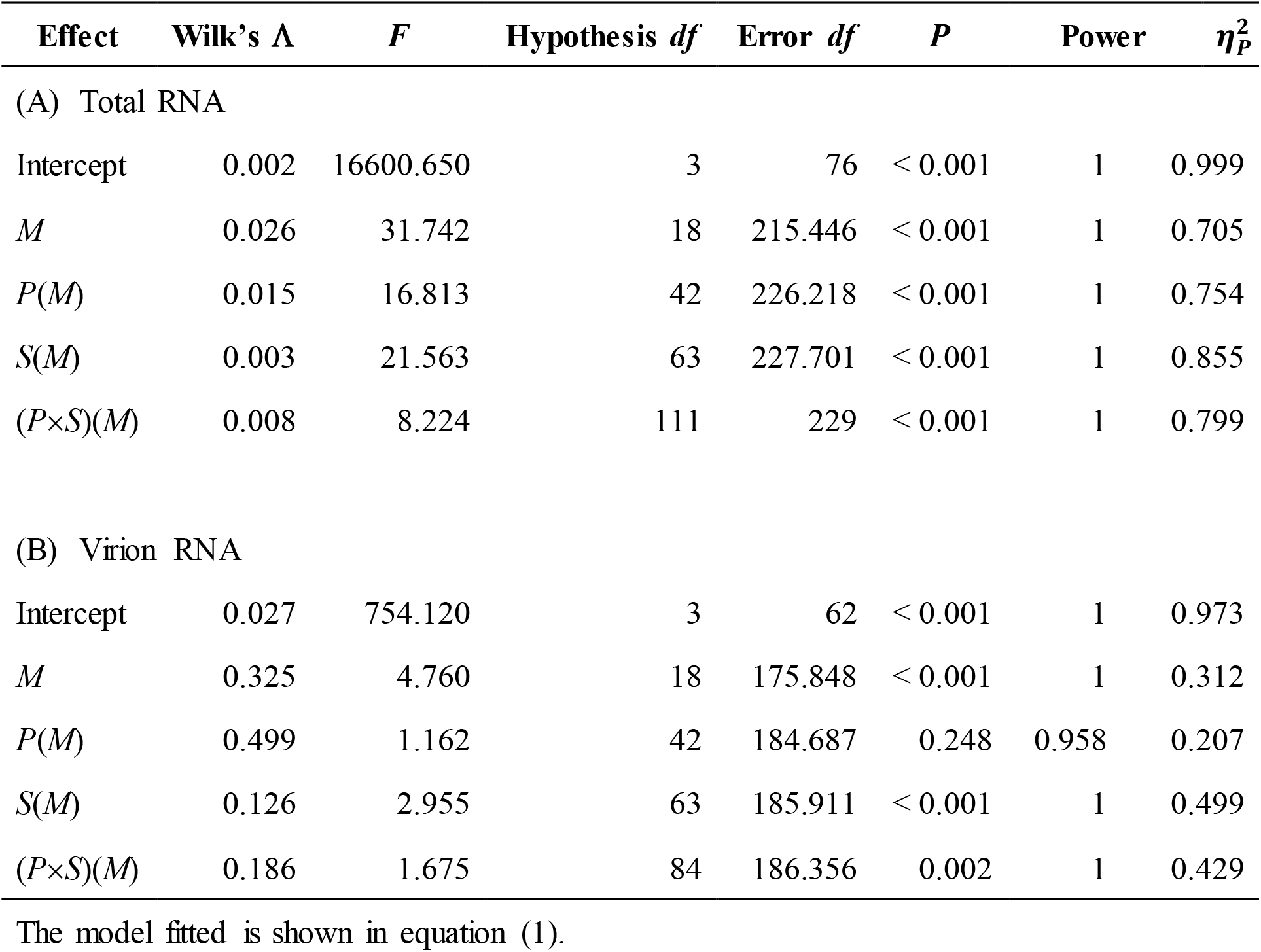
Results of the MANOVA analysis for the RNA frequencies estimated from total and from virion RNA extractions in experiments with variable input ratios done in *N. benthamiana*. The model fitted is shown in equation (1).

| Effect | Wilk's $\Lambda$ | $F$ | Hypothesis $df$ | Error $df$ | $P$ | Power | $\eta_p^2$ |
| --- | --- | --- | --- | --- | --- | --- | --- |
| (A) Total RNA |  |  |  |  |  |  |  |
| Intercept | 0.002 | 16600.650 | 3 | 76 | < 0.001 | 1 | 0.999 |
| $M$ | 0.026 | 31.742 | 18 | 215.446 | < 0.001 | 1 | 0.705 |
| $P(M)$ | 0.015 | 16.813 | 42 | 226.218 | < 0.001 | 1 | 0.754 |
| $S(M)$ | 0.003 | 21.563 | 63 | 227.701 | < 0.001 | 1 | 0.855 |
| $(P \times S)(M)$ | 0.008 | 8.224 | 111 | 229 | < 0.001 | 1 | 0.799 |
| (B) Virion RNA |  |  |  |  |  |  |  |
| Intercept | 0.027 | 754.120 | 3 | 62 | < 0.001 | 1 | 0.973 |
| $M$ | 0.325 | 4.760 | 18 | 175.848 | < 0.001 | 1 | 0.312 |
| $P(M)$ | 0.499 | 1.162 | 42 | 184.687 | 0.248 | 0.958 | 0.207 |
| $S(M)$ | 0.126 | 2.955 | 63 | 185.911 | < 0.001 | 1 | 0.499 |
| $(P \times S)(M)$ | 0.186 | 1.675 | 84 | 186.356 | 0.002 | 1 | 0.429 |

However, a significant *P*-value tells nothing about the *magnitude* of the effect that a factor has on the measured variables; a small effect may still be significant. To assess the magnitude of effects we used the 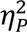 statistic that represents the proportion of total variability attributable to a given factor while controlling for all other factors. The advantage of 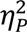 respect other measures of effect magnitude is that it allows for comparisons among different experimental designs. Conventionally, values of 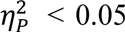 are considered as small, 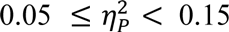 as medium and 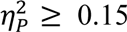 0.15 as large.Table1A shows that significant differences exist among plants inoculated with the same mixture and among equivalent samples from different plants and that the effect associated with these two factors is large (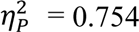 and 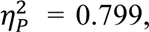 respectively). These differences appear as an unavoidable consequence of the stochastic events that take place during inoculation of different plants as well as during the progression of infection (*e.g.*, bottlenecks during cell-to-cell and systemic movement of viral particles [26]). Despite these differences, significant effects have been detected for the other factors.

The largest effect 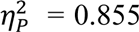= 0.855 is associated to differences among samples (*i.e.*, tissues). Differences among samples result from the fact that different tissues are infected with viral populations at different stages in their within-host evolution [27], with the lowest and oldest tissues containing viral populations at early stages of replication, whilst youngest tissues contain viral populations that have evolved longer and are closer to the equilibrium. The weakest effect, 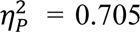, corresponds to differences among inoculation ratios, though it can still be considered as a very strong 192 effect.

The grand mean estimate for the relative frequencies corresponds to a *SGF_total_* of 1:3:2.Fig. 1A is a normalized ternary plot showing the marginal average of output ratios estimated for each input ratio. Regardless of the initial conditions, after infection all RNA populations tend to a rather limited region of the possible space of solutions, which contains the *SGF_total_* ratio.Table 1B shows the results from the MANOVA analysis run for the RNA frequencies estimated from encapsidated RNAs fitted to equation (1). The only difference with the results just reported for the total RNAs is the lack of differences among replica plants (factor *P*; *P* = 0.248). The magnitude of significant effects is 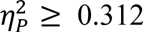, which is smaller than found for the case of total RNA but still large. In this case, the grand mean estimated for relative frequencies corresponds to a *SGF_encap_* of 1:2:1, a value that slightly differs from the one reported above for total RNAs, although in both cases shows that RNA2 is the most abundant one.Fig. 1B summarizes the evolution of segments ratio from the 206 input mixture to the average values obtained at the time of analyzing the encapsidated RNAs extracted. As above, values converge to a particular region of the space that 208 contains the *SGF_encap_* value.

**Fig. 1.**
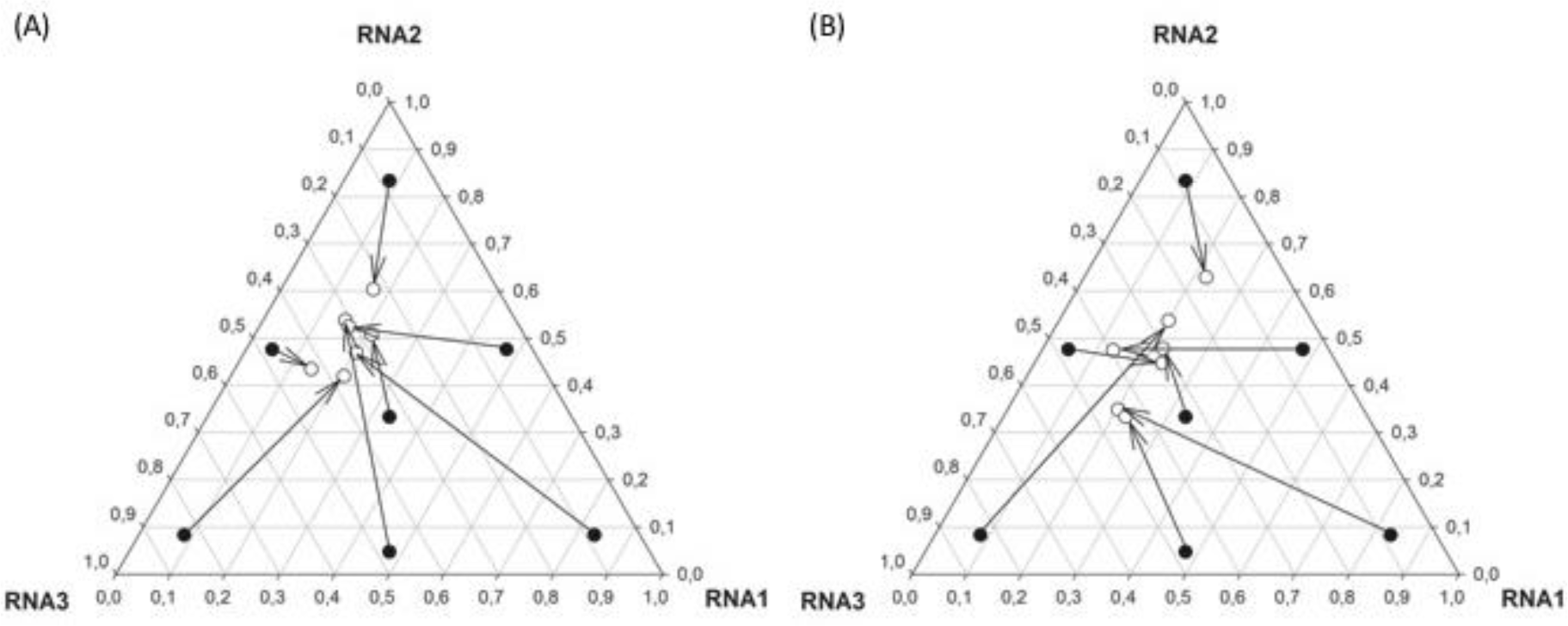
Normalized frequency ternary plot showing the abundance of each genomic RNA. Solid circles represent the inoculation rates. Open circles the marginal mean estimates of relative ratios at the end of the experiment for each input ratio. Arrows connect initial and final ratios.(A) Abundances estimated from total RNA samples.(B)Abundances estimated from RNA extracted from viral particles.

Overall, the results summarized in Fig. 1 suggest the existence of an attractor region in the frequencies phase diagram to which RNA populations converge after infection. In the following section we will explore whether (*i*) this equilibrium is driven by frequency-dependent selection (FDS) and (*ii*) it is stable [28].

### AMV *SGF* is driven by FDS to a stable equilibrium

Fig. 2 shows the graphical analysis of FDS as a driving force of the observed *SGF*. Marginal mean frequency data shown in Fig. 1 for each RNA segment were transformed into relative abundances as described in the Methods section and then log-transformed. The plots show the output log-relative abundances as a function of the input log-relative abundances.Fig. 2A shows the results for the RNA segments quantified in the total RNA samples. The dashed line represents the expectation under the null hypothesis of no-FDS [28]. For each RNA, we computed the output and input log-relative abundance and plotted them (different symbols). First, the data were fitted to a set of models with increasing number of parameters, but in all cases the linear regression was the best-fitting model (shown as solid lines). In all three cases, statistical significance of the FDS is tested by the deviation of the slope of the linear regression from 1 (the dashed diagonal). All three regression lines in Fig.1A have a slope significantly less than 1 (*t*_5_ ≥ 6.964, *P* ≤ 0.001). The analysis of Fig. 1A provides additional information of considerable biological interest, *i.e.*, whether the FDS is linear or not, how strong it is, and if an equilibrium point exists. A point of equilibrium occurs if and where the regression line crosses the diagonal [28]. At such point the focal RNA segment frequency is at the same frequency as expected in the absence of FDS. In all three cases, the slope is less than one, meaning that the equilibrium is evolutionarily stable; the system evolves towards an equilibrium *SGF_total_* that is stable against random perturbations of any of the three RNA components. For instance, perturbations may be associated with the inoculation process or by bottlenecks inherent to systemic movement and colonization of new growing tissues in the apical meristem [26].

**Fig. 2.**
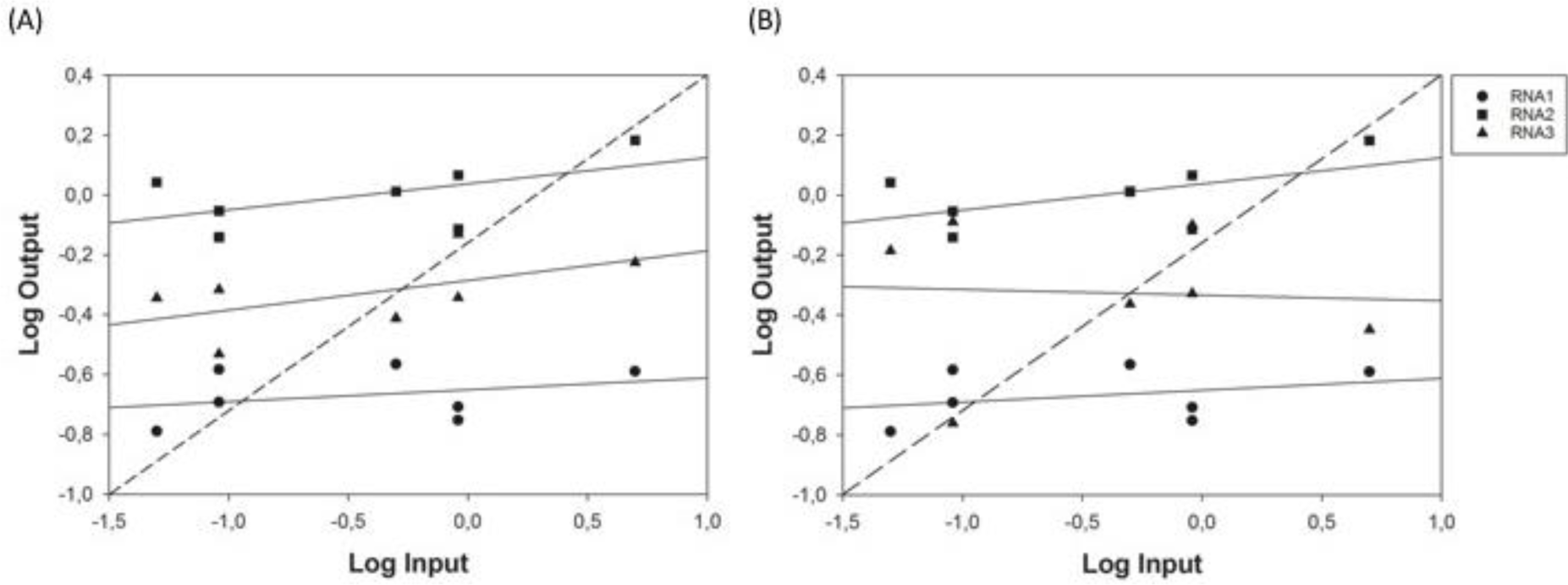
Graphical analysis of FDS as a mechanism to explain the relative abundance of the three RNA segments. (A)Abundances estimated from total RNA samples.(B) Abundances estimated from RNA extracted from viral particles. The dashed line corresponds to the null hypothesis of no-FDS. The continuous lines show the best fitting linear model to the relative abundances of each RNA.

Fig. 2B presents the graphical analyses of FDS for the encapsidated RNA ratios shown in Fig. 1B. The conclusions are qualitatively the same as those described in the previous paragraph for the total RNA samples: all three relationships are linear, with slopes significantly less than 1 (*t*_5_ ≥ 13.610, *P* ≤ 0.001) and thus the *SGF_encap_* also corresponds to an evolutionarily stable equilibrium.

To further characterize the nature of this FDS, we have analyzed the particular relationship between the marginal mean abundances of RNAs in both types of samples, from total and encapsidated RNAs. To do so, we have computed partial correlation coefficients among RNA abundances using as control variables the input ratios (*M* in equation (1)), plant replicate (*P* in equation (1)) and tissue sampled (*S* in equation (1)).Table 2 shows the hemi-matrix of correlations (notice that the matrix is symmetrical and thus the upper half has been removed). Focusing first in the quantifications from the total RNA extractions, we found that the synthesis of RNA3 negatively impacts the production of both RNAs 1 and 2, while the levels of RNAs 1 and 2 production do not affect each other. Looking now at the correlations between encapsidated RNAs, we found that all are negatively correlated with each other, thus suggesting that they compete for available capsids, which should then become a limiting factor. Finally, looking at correlations between total and encapsidated RNAs (non-gray cells in the hemi-matrix), with exception of RNA1, positive correlations exist between the amount of total and encapsidated RNAs (although the correlation for RNA3 becomes non-significant after accounting for multiple tests of the same hypothesis). RNAs 2 and 3 seem to strongly compete for encapsidation: the more RNA2 produced, the less RNA3 encapsidated and *vice versa*. However, RNA1 does not seem to be involved in this competition.

**Table 2.** Results from the partial correlation analyses among abundances of different RNA segments. Gray cells correspond to correlations among RNAs from the same type of sample. All tests have 79 *df*. Asterisks indicate cases significant after the Holm-Bonferroni correction of multiple tests of the same null hypothesis.

|  |  | Total RNA extraction |  |  | Virion RNA extraction |  |
| --- | --- | --- | --- | --- | --- | --- |
|  |  | RNA1 | RNA2 | RNA3 | RNA1 | RNA2 |
| **Total** | **RNA2** | $r = 0.024, P = 0.835$ | | | | |
| | **RNA3** | $r = -0.445, P < 0.001^*$ | $r = -0.895, P < 0.001^*$ | | | |
| **Virion** | **RNA1** | $r = -0.001, P = 0.991$ | $r = -0.049, P = 0.661$ | $r = 0.052, P = 0.645$ | | |
| | **RNA2** | $r = -0.021, P = 0.850$ | $r = 0.319, P = 0.004^*$ | $r = -0.282, P = 0.011^*$ | $r = -0.425, P < 0.001^*$ | |
| | **RNA3** | $r = -0.020, P = 0.863$ | $r = -0.278, P = 0.012^*$ | $r = -0.238, P = 0.032$ | $r = -0.418, P < 0.001^*$ | $r = -0.644, P < 0.001^*$ |

### AMV *SGF* varies among host species

Next we explored to which extent the host species determines the value of *SGF*. To do so, we inoculated five different susceptible hosts (*N. benthamiana*, *Nicotiana tabacum*, *Cucurbita pepo*, *Medicago sativa*, and *Capsicum annuum*) with a 1:1:1 mixture of the genomic RNA segments and evaluated the output frequency of each segment 7 dpi for *N. benthamiana* and 12 dpi for the rest of species, following the same sampling scheme than in the experiments previously described. Quantifications obtained for encapsidated RNAs from *M. sativa* and *C. annuum* were not reliably different from negative controls and thus were not considered for the following analyses. Frequency data were fitted to equation (2) in Methods using MANOVA and the results from this analysis are shown in Table 3. In case of segment frequencies in the total RNA extraction, all factors had a highly significant effect, with magnitudes being in all cases 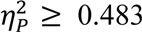 0.483 (Table 3A). There is great variation in the segments ratio among host species (Fig. 3), although the estimates for both *Nicotiana* spp. remain closer among them than they are relative to the other species analyzed. The grand mean value *SGF_total_* across hosts is 1:3:12, a value that sharply contrasts to the above stable equilibrium value found for *N. benthamiana* (1:3:2) due to the larger amount of RNA3 in this later case. In case of segment frequencies in the encapsidated RNAs, the only not significant factor was the interaction between plant replicate and type of sample ((*P* × *S*)(*E*) in equation Table 3B), although highly significant differences exist among host species. In this case, the grand mean value *SGF_encap_* across hosts is 1:3:2, which also differs from the 285 stable equilibrium value found for *N. benthamiana* but to a lesser extent.Fig. 3 also shows that estimates obtained from total and encapsidated RNA extractions render values that are close in the normalized ternary plot, thus showing a good correlation among them. Therefore, we can conclude that the segments ratio at 12 dpi strongly depends on the host species which is being infected, which suggests the *SGF* is host-species dependent.

**Fig. 3.**
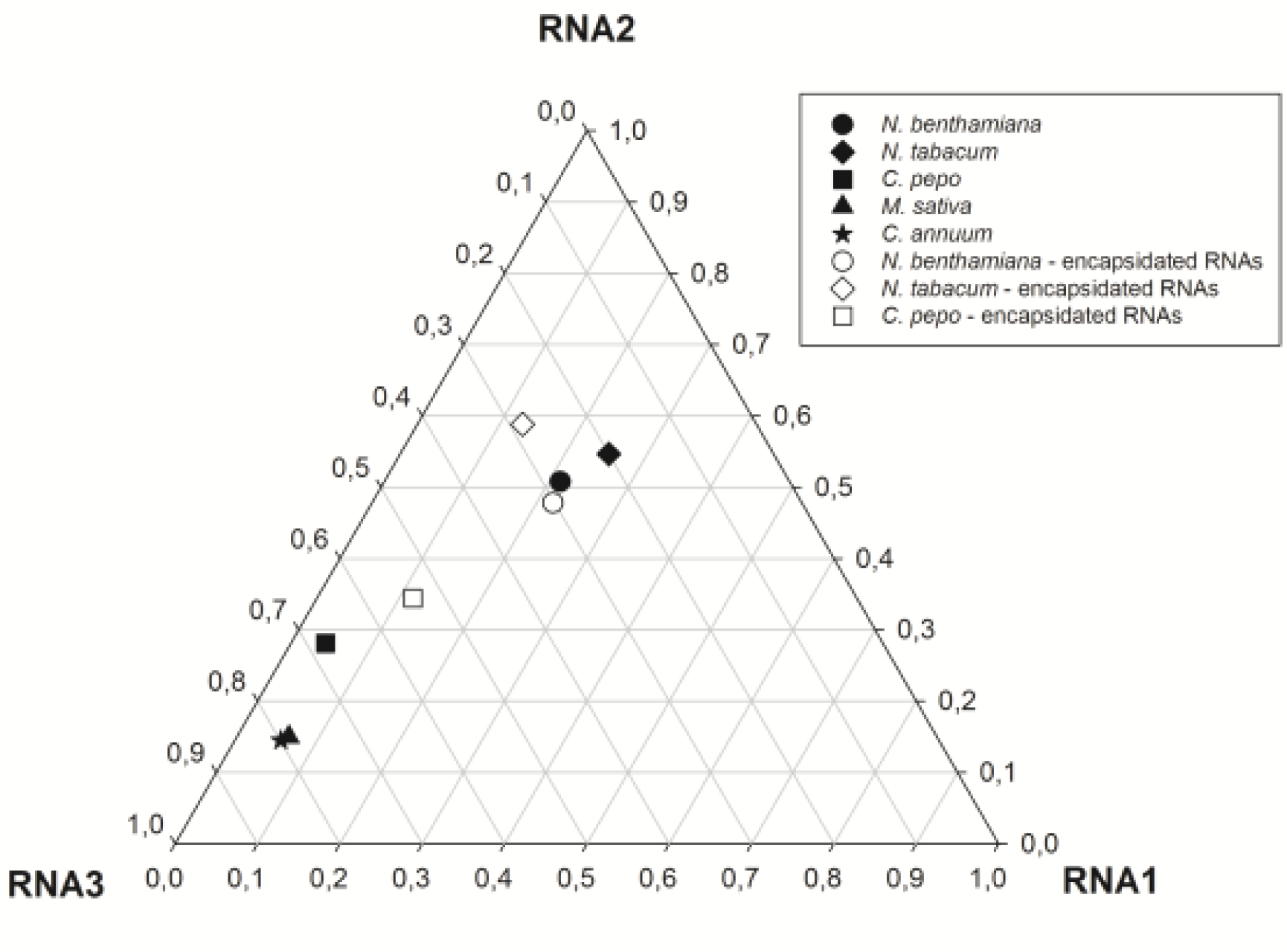
Normalized frequency ternary plot showing the abundance of each genomic RNA on different plant hosts. Solid symbols represent the marginal mean frequencies estimated from total RNA samples. Open symbols represent the marginal mean frequencies estimated from virion RNA samples. In the latter, no data are available for *C. annuum* and *M. sativa*.

### Total RNA production is maximized at the *SGF_total_*

We have observed that *SGF_total_* is maintained by a *FDS* mechanism, and that the actual value taken by the *SGF_total_* depends on the host wherein AMV replicates. Has *SGF_total_* been optimized in each host by natural selection to maximize the total accumulation of the three genomic RNAs? To tackle this question, we have computed a partial correlation coefficient between the total RNA accumulation (summing up the accumulations of the three RNA segments) and the Euclidean distance computed between the *SGF* values obtained for each individual sample (*i.e*., a particular tissue from a given plant from each host species) using as control 301 variables the host species (*E* in equation (2)), input ratios (*M* in equation (1)), plant replicate (*P* in equation (2)) and tissue sampled (*S* in equation (2)). The rationale behind this test is as follows: if *SGF_total_* has been optimized to maximize the 304 production of AMV genomic RNAs, then the farther from the evolutionarily stable *SGF_total_* equilibrium the replicating AMV population, the lower the accumulation of genomic RNAs. Conversely, the closer to the equilibrium *SGF_total_* a replicating viral population would be, the higher the accumulation of AMV RNAs. A low yet highly significant negative correlation exists between distance to the optimal *SGF_total_* and total RNA accumulation (*r* = −0.249, 120 d.f., 1-tailed *P* = 0.003), thus backing up the 310 hypothesis that RNA accumulation is maximal at the equilibrium *SFG_total_*.

By contrast, no significant correlation exists between the *SGF_encap_* and the total amount of the three RNAs encapsidated (*r* = −0.133, 89 d.f., 1-tailed *P* = 0.105), suggesting that the strength of selection for encapsidation has been weaker than for replication.

## Discussion

Here, we have explored the within-host evolution of the ratio of the three genomic segments of the multipartite plant virus AMV. We found that regardless the ratio used at inoculation, an evolutionary stable equilibrium is reached in which 321 the three RNA segments are represented in a 1:3:2 stoichiometric ratio, in *N. benthamiana*. The occurrence of a stable genome-segment formula has been dubbed as the setpoint genome formula, or *SGF*, by Sicard et al. [15], who described within-host evolution towards a stable composition for a multipartite DNA virus, FBNSV. Here, we have extended this observation to a second multipartite virus, which is more representative of the vast majority of multipartite viruses by virtue of being an RNA virus. We have observed that the ratio of encapsidated segments also represents an evolutionary stable equilibrium but with a slightly different solution: 1:2:1. Related to these observations, we found that the production and encapsidation of the different segments are linked in a non-trivial manner. We speculate that 331 RNAs 1 and 2 cooperated during replication whilst competing with RNA3, and that all three RNAs competed with each other for the CP for encapsidation. Although these inferences on the interactions between the genome segments are based on a correlation analyses, they do reflect a biologically relevant association between viral traits. However, we have not tested the underlying mechanisms, and do not know whether there is a direct causal link between both traits or the correlation is mediated by a third yet unknown factor. Future work will explore the mechanisms of these 338 correlations.

We found that the *SGF* appears to be dependent on the host species, suggesting the involvement of host factors that differnt among host species play a role in its regulation. In agreement with our findings, Ni et al. [29] also found that the relative abundances of encapsidated and total RNA segments of BMV were also dependent on the host species. Indeed, when the ratios of the three segments were followed during the progression of infection in two monocot hosts, they converge into stable *SGF_total_* (1:2:3 for barley and 1:2:2 for wheat). However, at odds with our findings, these authors concluded that no relationship existed between the total RNAs produced and their relative encapsidation. Nonetheless, ignoring the fact that the percentages 348 reported in their Fig. 1 not always add up to 100%, a significant correlation exist between the relative frequencies of total and encapsidated RNAs in wheat (Spearman’s *ρ* = 1, 2 d.f., *P* < 0.001) but not in barley (*ρ* = 0.400, 2 d.f., *P* = 0.600).

Multiple theories have been proposed to explain the existence of multipartite viruses, most commonly found in plants. The most recent and tantalizing proposal is that genome segmentation represents an efficient and rapidly adaptable way of 354 regulating gene expression throughout manipulation of gene copy numbers [15]. As the changes observed here in the *SGF* in alternative host species occurred within a narrow time window, multipartition might be advantageous for rapid adaptation of gene expression in a manner that is largely nucleotide-sequence independent, and therefore also mutation independent. Such an approach to adaptation could be especially advantageous in alternative hosts, where founder numbers may be small due to low infection probabilities and effective population sizes might also not be large initially due to poor replication.

On the other hand, according to this hypothesis, a tight link must exist between the necessity of producing a given protein and the abundance of the RNA segment that encodes for it. At first glance, this hypothesis does not apply to AMV for two reasons. Firstly, as one may imagine that CP, necessary for producing infectious virions and encoded by the RNA3, would be required in larger numbers than the replicase complex, encoded by RNAs 1 and 2. However, it is important to recall at this point that CP is translated from a subgenomic RNA. We have not quantified the abundance of RNA4. Interestingly, however, we observed that the ratio of RNAs 1 370 and 2 remains more or less constant in all experimental conditions tested (see below). Secondly, our observation of RNA2 accumulating more than RNA1 may suggest that P2 should also accumulate more than P1. Unfortunately, no quantitative data are available on the accumulation of AMV P1 and P2 in virus infected tissues. Comparing with other members of the *Bromoviridae* family, it has been shown that BMV and *Cucumber mosaic virus* (CMV) 1a protein accumulates to larger amounts than the corresponding 2a protein in purified replication complexes [30-32], suggesting that *trans* elements are controlling the translation of viral RNAs. In this sense, it has been observed that AMV CP enhances the translational efficiently of viral RNAs *in vivo* [33] via the interaction with the 3’ termini, which adopts two alternative structures for translation (a linear array of hairpins with high affinity for CP) and replication (a pseudo-knotted structure) [34]. A similar mechanism has been reported as regulator for translation of the replication complex proteins of BMV [35]. The assumption that the amount of protein expressed is always proportional to the amount of messenger RNA, although appealing, has been proven wrong. For example, during mixed phage infections of bacterial cells, increasing the number of genomic copies results in switches between lytic and lysogenic states and the concomitant production of viral proteins [36]. Indeed, in such instances the 388 regulation of gene expression is an emerging property of the structure of regulatory networks rather than directly resulting from gene copy number [36]. Translation efficiency and RNA stability are inexorably linked [37], further challenging simple interpretations of the effects of observed RNA levels on actual protein expression levels.

We found significant differences between the *SGF* estimated from total RNA production and from encapsidated RNA. Selection may operate in distinct ways here, eventually resulting in an evolutionary tradeoff. On the one hand, within-host selection on replication will result in a *SGF_total_* that maximizes replication of the three RNA segments, likely by producing an optimal combination of RNAs and proteins (*i.e.,* regulation of gene expression), thus a segments ratio that would necessarily depart from the 1:1:1 as more proteins encoded by one segment are needed than proteins from other segments. This possibility is clearly supported by the negative correlation that we have observed between the total amount of genomic RNA produced and the distance to the *SGF_total_*. On the other hand, selection operating at 403 the between-host level will result in a *SGF_encap_* that maximizes the probability of a successful transmission, thus closer to the 1:1:1 ratio. Our observations do not back up this possibility, as we have not observed the predicted negative correlation between total encapsidated RNAs and the *SGF_encap_*. This negative correlation between within-host accumulation of viral RNA segments and the amount of each segment encapsidated and available for transmission is linked to the classic tradeoffs between within-host growth and between-host transmission brought forward to explain the virulence of vector-borne pathogens [38]. According to this tradeoff, virulence is an unavoidable consequence of within-host multiplication of parasites and thus within-host selection would result in increases in accumulation and thus in virulence. However, virulence reduces the chances of the pathogen to be transmitted and thus virulence should evolve to an intermediate value that maximizes its transmission and thus its *R*0 fitness value. Our results support this possibility, as selection has improved within-host accumulation without enhancing transmission probability. Very few studies have directly tackled the association between accumulation, virulence and transmission. In one of such study performed with the tripartite CMV, the evolution of tolerance mechanisms by *Arabidopsis thaliana* allowed for high accumulation with low virulence [39], thus rejecting the hypothesis. In another study, the expected correlations between accumulation and virulence and between virulence and transmission were observed for the monopartite *Cauliflower mosaic virus* [40].

The mechanisms that may determine AMV’s *SGF* remain elusive. Probably the evolutionarily stable *SGF* results from complex molecular interactions between viral and host components, inextricably intertwined with viral population dynamics. Some evidences available in the literature may help to bring light into this complex question. For instance, results from transient expression experiments of proteins P1 and/or P2 revealed that replication of RNAs 1 and 2 depends on the presence of these proteins in *cis* and that, within infected cells, the replication of RNAs 1 and 2 is strictly coordinated through the encoded proteins rather than by RNA-RNA interactions [41]. This coordination may ensure the expression of proteins P1 and P2 in the correct ratio to form the replication complex. However, the replication of RNA 3 is not linked to the replication of RNAs 1 and 2 [42, 43]. In this sense, these interactions explain the results reported in Table 2, namely, the negative correlation observed between replication of RNA3 and production of RNAs 1 and 2: replication of both RNAs 1 and 2 is coupled and not interfering each other, while the replication of RNA 3 must use the full replicase complex in *trans* and, thus, competes with the replication of RNAs 1 and 2. These observations lead to the prediction that the ratio between RNAs 1 and 2 could be constant (see below) but also suggest that the different RNAs of segmented viruses could be considered as independent molecules unless they replicate coordinately, with *cis* elements that constraint the accumulation of the corresponding viral RNAs. It is tempting to speculate that virus resistance mediated by the expression of viral proteins – normally the CP in transgenic plants – could be at least partially due to a drastic alteration of the genome segment ratio and the negative effects thereof on viral replication.

Regarding the results obtained from hosts other than *N. benthamiana*, first we must acknowledge a limitation of our experimental design: As we did not consider different starting ratios or multiple time points post inoculation, one could question whether virus populations have reached a stable *SGF*. Conservatively speaking, we can only conclude that at advanced stages of infections in all hosts (*i.e*., 12 dpi), the ratio of segments significantly differs among hosts and significantly departs from the one value estimated for *N. benthamiana*. This being said, we observed that the ratio between RNAs 1 and 2 remains constant (ca. 1 RNA1 molecule per 3 RNA2 molecules) probably due to the coordinated replications between both RNAs, as mentioned in the previous paragraph. The coordinated replication of both RNAs 1 and 2 may determine the ratio of both viral RNAs independently of the host species. In this sense, we observed that the ratio of both viral RNAs oscillated between 1:2 - 1:3. Apparently, the system dynamically evolves to maintain the correct ratio between RNAs 1 and 2, to the detriment of the accumulation of the RNA 3. Furthermore, the accumulation of RNA 3 was significantly altered depending of the host species, indicating that the virus may use this RNA to accommodate its life cycle to the presence/absence of different host factors, for instance, the transcription factor promoting salicylic-dependent defense signaling response recently reported to interact with it [44].

All three AMV RNAs contain binding sites for the CP at the 3’UTR and bind it with an equal distribution between all viral RNAs [45, 46]. In solution, AMV CP occurs as dimers and these dimers are the building blocks of viral capsids [45]. N-terminal peptides of CP bind to the 39 nucleotides of the 3’UTR RNAs in a 2:1 stoichiometric ratio [47]. Binding of the CP to the 3’UTR also enhances translation of viral RNAs by mimicking the function of the host poly(A)-binding protein [48, 49]. Altogether, these evidences point to the idea of a CP with multiple functions that are critical at different steps of the virus infectious cycle. The results obtained in the present work support the idea that the RNAs are competing for the CP, and it is therefore a limiting factor that could be used for interventions aimed at controlling virus infection. In agreement with this result, it has been recently observed that AMV CP accumulated at the nucleus and nucleolus, an observation interpreted as a mechanism to control virus expression by the cytoplasmic/nuclear balance of CP accumulation [50].

## Methods

### Host species and virus inoculation

Plants from the experimental hosts *C. annuum* L., *C. pepo* L., *N. benthamiana* Domin, *N. tabacum* L. cv. Samsun, and *M. sativa* L. were all mechanically inoculated with a mixture of 5’ capped transcripts corresponding to AMV strain RNAs 1, 2 487 and 3 plus a few μg of purified AMV CP as described previously [51]. For the transcription reactions, clones pUT17A, pUT27A and pAL3-NcoP3, containing full-length cDNAs of AMV RNAs 1, 2 and 3, respectively, were linearized with appropriate restriction enzymes and transcribed with mMESSAGE mMACHINE^®^ T7 kit (Ambion, USA). The quantification of the AMV RNAs was performed with a ND-1000 spectrophotometer (Thermo Scientific, USA) and agarose gel eletrophoresis using an RNA ladder (RiboRuler High Range RNA Ladder 200 to 6000, Thermo Scientific) and several dilutions of the transcribed RNAs.

Before addressing the specific questions of this study, we estimated the minimal amount of AMV transcripts required to initiate an infection in the different hosts by performing serial dilutions of an initial inoculum mixture with a ratio 1:1:1. Henceforth, all ratios of AMV genomic RNA segments are given as RNA1:RNA2:RNA3. For *N. benthamiana* plants we selected a final concentration of each AMV RNAs of 40 ng/μL each, whereas for the rest of hosts it was necessary to increment the transcripts concentration five times (200 ng/μL each RNA).

All species were inoculated with the AMV RNAs ratio of 1:1:1 (three plants per ratio) except *N. benthamiana* plants that were also inoculated with ratios: 10:1:1, 504 1:10:1, 1:1:10, 10:10:1, 10:1:10, and 1:10:10. For each of these experiments, at least three plants were inoculated. All plants were grown in a biosafety level-2 greenhouse at 24/20 °C day/night temperature with 16 h light. After 7 (*N. benthamiana*) or 12 (rest of species) days post inoculation days (dpi), all inoculated plants were analyzed for the abundance of each RNA segment in both total RNA extraction and virus particle purification from inoculated, middle and upper leaves and from the remaining tissues of the plants (*i.e*, four samples per plant).

### Virus particles purification and total RNA extraction

Leaves or plants were homogenized with mortar and pestle in liquid N2 to minimize the putative irregular virus distribution in the tissue. Total RNA extraction was performed using 0.1 g of tissue and the Plant RNA Isolation Mini Kit (Agilent, USA) following the manufacturer’s protocol. All samples were diluted to a final concentration of 50 ng of total RNA/µL. Virus particles purification was performed using 0.5 g of the homogenized tissue, following the protocol previously described [52]. The fraction of enriched virus particles was resuspended in 100 μL of PE buffer (10 mM NaH2PO4, 1 mM EDTA, pH 7.0), that was subsequently subjected to RNA extraction using the Plant RNA Isolation Mini Kit (Agilent, USA). All RNA samples were stored at −80 °C until use.

### Quantification of AMV RNAs by RT-qPCR

The standard curves to quantify the AMV RNAs 1, 2 and 3 in the samples by RT-qPCR were prepared using known amounts of DNase-treated transcripts derived from the linearized pUT17A, pUT27A and pAL3-NcoP3 plasmids. To ensure a correct estimation of the transcripts concentration, all sample were analyzed with a ND-1000 spectrophotometer (Thermo Scientific, USA) and by agarose gel electrophoresis. To construct the standard curve for each RNA, we selected six (RNAs 2 and 3) or seven (RNA 1) different viral RNAs concentrations, calculated in terms of copy number molecules per microliter (www.endmemo.com/bio/dnacopynum.php), that were generated by 5-fold serial dilutions of a starting solution containing 10^10^ (RNA 1) or 2×10^9^ (RNAs 2 and 3)molecules of the corresponding viral RNA per μL. All dilutions were made in a solution containing 50 ng/μL of total RNA extracted from healthy *N. benthamiana* plants.

The primers used for amplifying RNAs 1, 2 and 3 were designed using PrimerQuest^®^ Design Tool version 2.2.3 (IDT Inc., USA), selecting the parameters GC% = 40 - 60%, Tm = 57 - 60 °C, and size = 100 - 150 bp. The primers used for the RT-qPCR reactions for AMV RNAs 1, 2 and 3 are listed in Supplementary Table S1. To estimate the number of genome equivalents present and their frequencies, all data for the standard curve were first log-transformed to ascertain the range over which the response was linear. The dynamic range was limited to one dilution before the response appeared to saturate. Linear regression of the log-transformed data was then performed, rendering high values for the determination coefficient (*R*^2^ >0.98) and of the slope-derived amplification efficiency (90% - 110%). For those samples that fell within the dynamic range, the estimated linear regression parameters were used to estimate the unknown concentrations in the virus samples.

Duplicated RT-qPCR reactions were carried out in 10 μL reaction volume using the GoTaq^®^ 1-step RT-qPCR system (SYBR^®^ Green) (Promega, USA) and the StepOnePlus Real-Time PCR System (Applied Biosystems, USA). Each reaction contained 50 ng RNA sample, 5 μL of the 2 ×master mix, 10 μM of both the forward and reverse primer, 0.2 μL of GoScript^TM^ RT Enzyme Mix and 0.155 μL of CXR reference Dye (30 μM). The reactions were incubated at 42 °C for 15 min, followed by 95 °C during 10 min and 40 cycles of 95 °C for 10 s, 62 °C for 34 s and 72 °C for 30 s. After the RT-qPCR reaction, the melting curve stage was determined by incubating 95 °C for 15 s, 60 °C for 1 min and 95 °C for 15 s. The quantification of RNAs 1, 2 and 3 copy number was calculated using the StepOne Software v.2.2.2 (Applied Biosystems, USA).

Supplementary File S2 contains the absolute quantifications of the three RNA segments for all the experimental samples used in this study.

### Statistical methods

The number of copies of RNA segment *i*, *Ri*, on each sample were transformed into relative frequencies, *f_i_*, by dividing them by the sum of the values estimated for every RNA segment on the corresponding sample, averaged across the two technical replicates of RT-qPCR: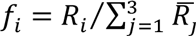. To analyze the effect that differen inocula mixtures of the three RNA segments had on the outcome of infection, frequency data were fitted to a multivariate linear model using MANOVA techniques. The model equation fitted reads:

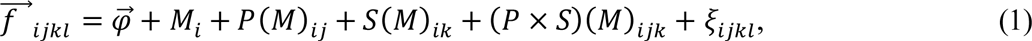

where 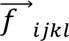 is the vector of frequencies measured for technical replicate *l* ∈ {1,2} of sample *Sk* (*k* ∈ {inoculated leaf, middle leaf, upper leaf, the rest of the plant (stems + apical tissues)}) taken from plant replicate *P_j_* (*j* ∈ {1,2,3}) that was inoculated with a mixture *M_i_* (*i* ∈ {1:1:1, 10:1:1, 1:10:1, 1:1:10, 10:10:1, 10:1:10, 1:10:10}) of RNA segments. Factors *P* and *S*, as well as their interaction, were treated as orthogonal,and nested within factor *M*. *ξ_ijkl_* measures the experimental error and was assumed to be normally distributed.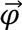 is the vector of grand mean frequency values and represents a statistical estimate of the *SGF*. Wilk’s Λ distribution was used for the multivariate tests of each factor in the model.

To analyze whether *SGF* depends on the host species inoculated with a 1:1:1 583 mixture, the corresponding frequency data were fitted to the following multivariate 584 linear model:

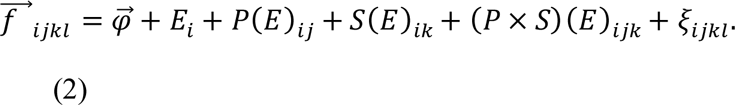

In this case, factor *Ei* represents the plant species (*i* ∈ {*C. annuum*, *C. pepo*, *M. sativa*,*N. benthamiana*, *N. tabacum*} and all other factors are as described for equation (1).

Next, we considered whether there was frequency-dependent evolution of the ratio of RNA segments infection of plants. In other words, we considered whether the frequency of one segment depends in a positive or negative manner on the abundance of the other two segments. Here, we made use of the classic population genetic approach described in [28]. In short, the ratio of the *j*^th^ RNA segment to its two counterparts was computed as Ω_*j*_ = *R_j_*/∑_*k*≠*j*_ *R_k_* for both the input mixture and the observed output mixture. In the absence of frequency-dependent selection (FDS),it is expected that the regression of the output 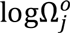 on the input 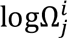 would be linear with slope one [28]. Significant deviations from the slope one relationship are taken as evidence of positive or negative FDS.

If FDS exists, then it can be evaluated whether (*i*) one or more equilibrium points exist and (*ii*) their stability. If the relationship between 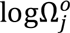 and 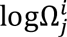 is linear, a single equilibrium point exists. If the relationship is not linear, then the number of equilibria equals the number of times the best-fitting function intersects with the diagonal of the 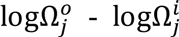 phase diagram (*i.e.*, the equation of slope one and intercept zero). Equilibria stability can be assessed by evaluating the value of the derivative 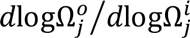 at the corresponding equilibrium point. 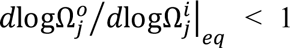 corresponds to an stable equilibrium in which the three segments coexist whereas 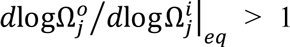 non-stable one in which the abundances of the three segments may experience changes due to very small perturbations.

In all cases, segment frequency data obtained from total RNA extractions and from virus preparations were analyzed separately. MANOVA and other statistical analyses were done using IBM SPSS version 23 (Armonk, NY, USA).

## Acknowledgements

This project was funded in part by grants BFU2015-65037P from the Spanish Ministry of Economy and Competitiveness, PROMETEOII/2014/021 from Generalitat Valenciana and EvoEvo (ICT610427) from the European Commission 7^th^ Framework Program to S.F.E. The China Scholarship Council and the Chinese Academy of Agricultural Sciences provided funding to B.W. We thank Francisca de la Iglesia, Paula Agudo and Lorena Corachan for their dedication and expert technicalassistance.

## Supporting Information

**S1 Table. Primers used to amplify each RNA segment.**

**S2 File. Excel file containing the average absolute quantifications for each 794 segment on each experimental sample.**

**Supplementary Table S1.**
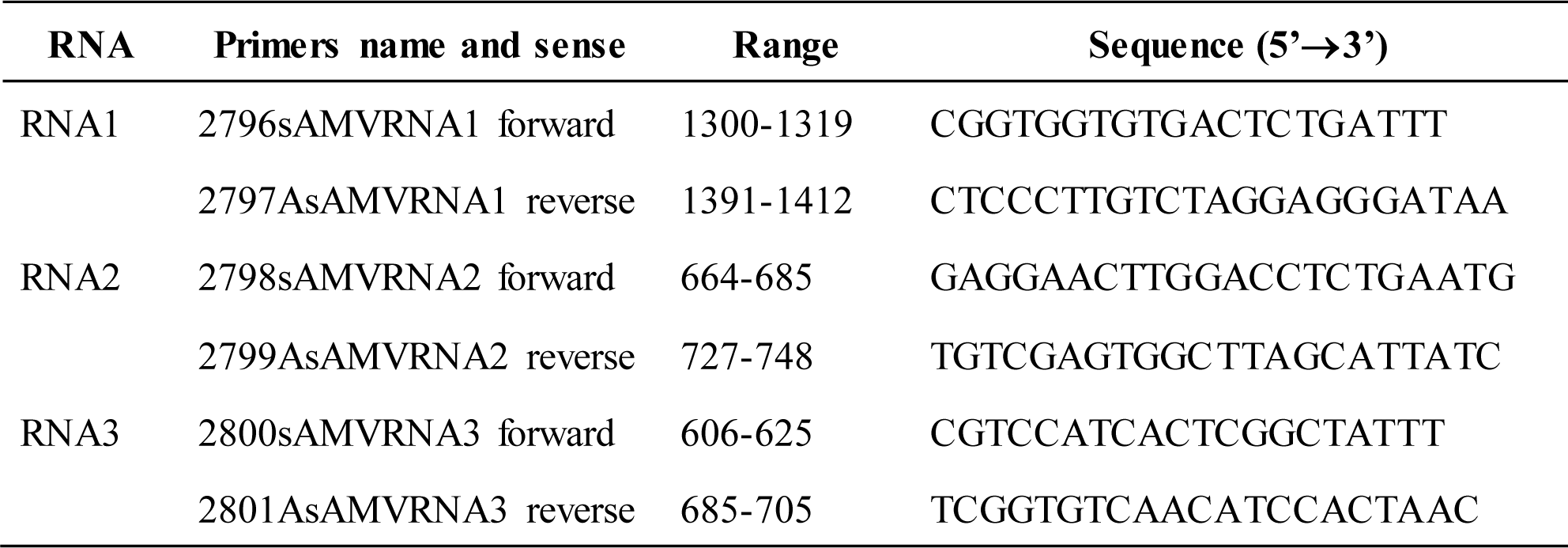
Primers used to amplify each RNA segment.

